# Single molecule targeted sequencing for cancer gene mutation detection

**DOI:** 10.1101/029686

**Authors:** Yan Gao, Liwei Deng, Qin Yan, Yongqian Gao, Zengding Wu, Jinsen Cai, Daorui Ji, Gailing Li, Ping Wu, Huan Jin, Luyang Zhao, Song Liu, Michael W. Deem, Jiankui He

## Abstract

With the rapid decline cost of sequencing, it is now clinically affordable to examine multiple genes in a single disease-targeted test using next generation sequencing. Current targeted sequencing methods require a separate step of targeted capture enrichment during sample preparation before sequencing, and the library preparation process is labor intensive and time consuming. Here, we introduced an amplification-free Single Molecule Targeted Sequencing (SMTS) technology, which combined targeted capture and sequencing in one step. We demonstrated that this technology can detect low-frequency mutations of cancer genes. SMTS has several advantages, namely that it requires little sample preparation and avoids biases and errors introduced by PCR reaction. SMTS can be applied in cancer gene mutation detection, inherited condition screening and noninvasive prenatal diagnosis.

## Introduction

In the past few years, the cost of large-scale DNA sequencing has been dramatically driven down by the tremendous advances in next-generation sequencing (NGS)^1^. Nonetheless, the cost of human whole genome sequencing and bioinformatics interpretation is still significant. In clinical practice, NGS is used to examine specific gene panels such as cancer genes and inherited conditions, sample numbers are high and data volume per sample is relatively small. It is often more cost-effective and time-efficient to target, capture, and sequence only the genomic regions of interest^2^. For example, there are several cancer gene panels commercially available, targeting as few as 50 to many hundreds of genes that are frequently mutated in cancer patients^3^. The cancer gene panel targeted sequencing has been proved to be useful in hereditary cancers diagnosis, and disease management.

Current NGS based targeted sequencing methods require a separate step of capture enrichment during sample preparation before sequencing^4, 5^. The two most commonly used custom-capture methods are based on hybridization or on highly multiplexed PCR. In the solution-based hybridization method, biotinylated DNA or RNA complementary probes are designed bind to gene targets, which are then purified using streptavidin-labeled magnetic beads. In the multiplexed PCR method, hundreds or thousands of PCR primer pairs are mixed to amplify the targeted genes.

In this report, we demonstrated a technology and platform to perform Single Molecule Targeted Sequencing (SMTS), which combined targeted capture and sequencing in one step. We used a combination of Total Internal Reflection Fluorescence (TIRF) microscope and single molecule fluorescence dyes to reject unwanted background noise and get single molecule resolution images^6^. The gene-specific flow cell was constructed with capture primers for gene regions of interest and the target genes can thus be sequenced without copying the DNA or enrichment before sequencing. Compared to current targeted sequencing methods with separate capture steps, SMTS has significant advantages, including little sample preparation and avoidance of biases and errors introduced by PCR amplification^7^. SMTS can be applied in cancer gene mutation detection, inherited condition screening, and high-resolution human leukocyte antigen (HLA) typing.

## Results

### Single molecule detection

The fundamental limitation of detection of single molecule fluorescence signals stems from the intrinsic qualities of the fluorophore. The key challenge is to reduce the background interference, which may arise from Raleigh scattering, Raman scattering, and contaminant fluorescence. Various single-molecule fluorescence microscopy techniques have been developed in the last two decades to overcome the difficulty in detecting single molecules with high signal to noise ratios in the presence of optical background^8^.

We applied Total Internal Reflection Fluorescence (TIRF) microscopy in this study. The optical setup is shown in Fig. 1. When light strikes an interface going from coverslip glass to fluid in the flow cell chamber at an angle greater than a critical angle, it undergoes a total internal reflection. This generates an exponentially decaying light field called the “evanescent wave” above the surface of glass. The evanescent wave excites fluorescent molecules within about 150-200 nanometers of the surface. The fluorescence from the labeled DNA molecules anchored on the glass surface is detected through a microscope objective and fluorescence filters by high sensitivity Electron-Multiplying CCD (EMCCD) cameras. As only the vicinity of the surface is illuminated, the noise from the bulk fluids of flow cell chamber is dramatically reduced. Single DNA molecules anchored on the surface can thus be monitored with high signal to noise (Fig. S1, S2 and S3).

**Figure 1.**
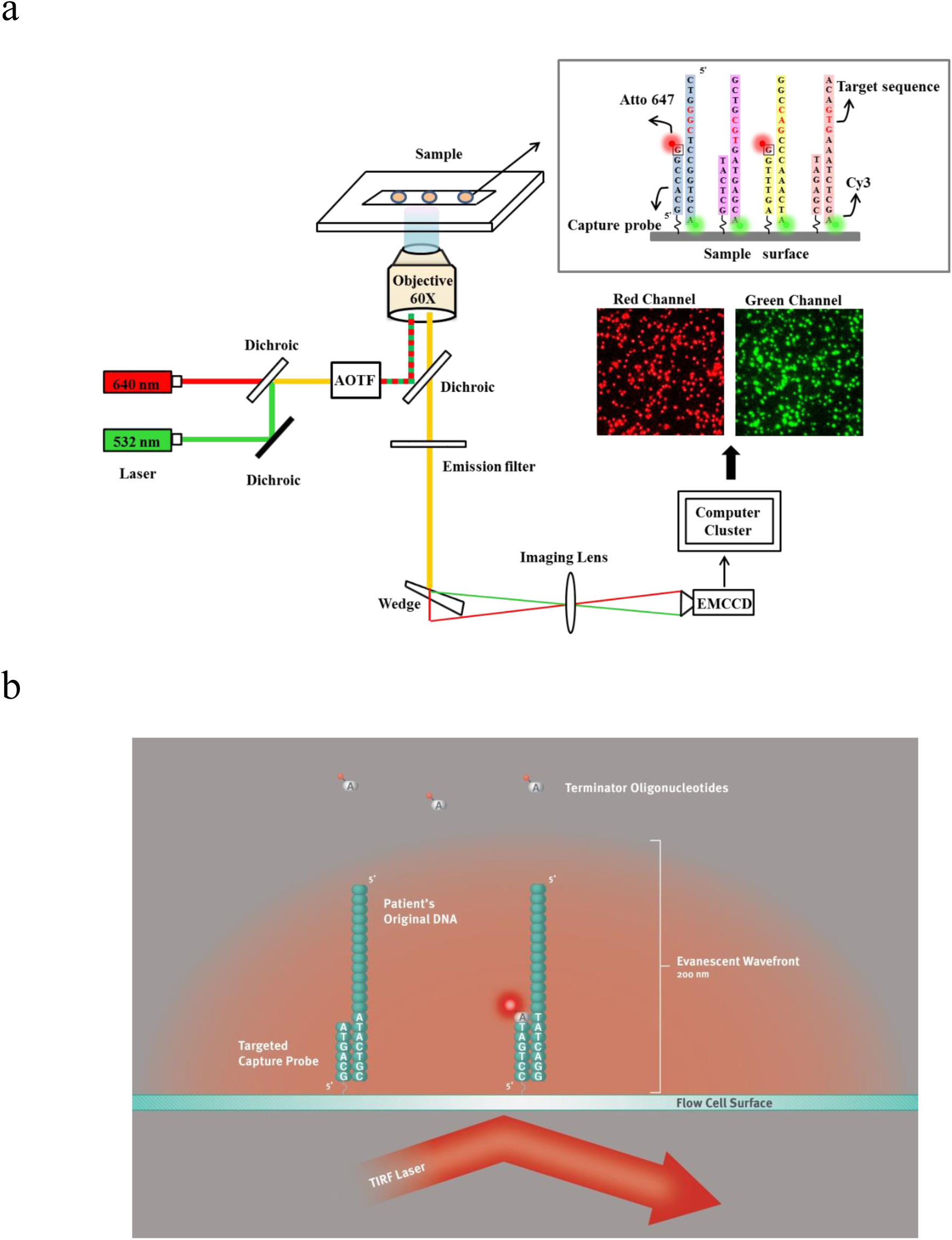
Schematic drawing of single molecule sequencing platform. (a) Schematic drawing of the optical setup. The green laser illuminates the Cy3 dyes which are attached to 3’ end of the target DNA template. The Cy3 dyes are non-cleavable. The red laser illuminates the cleavable Atto647N dyes which are attached to reversible terminators. Both Cy3 and Atto647N fluorescence spectra are recorded independently by an EMCCD. (b) Schematic of primed DNA templates attached to epoxy coated coverslip surface. The capture probes are covalently attached to the coverslip surface, and the target DNA templates are hybridized to the capture probes. The evanescent wave of TIRF illuminated the area within 200nm above the flow cell surfaces. The DNAs attached to the surfaces are within the range of evanescent wave.

The choice of fluorescent dyes to label nucleotides is also critical for single molecule detection. Many common fluorescent labels show rather low photostability if high-intensity laser excitation is used and processes are to be observed over long periods of time. We choose the ATTO 647N dyes to label the nucleotides, which fluoresces twice as strong as cyanine 5 in aqueous solution. Meanwhile, we optimized the imaging buffer to increase the photostability up to five times (Fig. S5).

Single-step photobleaching is used as a quality control to distinguish single molecule from multiple molecules. In an ideal situation, each DNA molecule is separately binding to the flow cell surfaces and the minimal distance between two DNA molecules is larger than the diffraction limit of light. In a random attachment cenari(as used in the present study) drive by Poisson statistics, two or more DNA molecules may bind to the surface at a distance less than the Rayleigh criterion. We quantified the amount of single DNA molecules to aggregated DNA molecules binding to the surface by observing the photobleaching patterns. The single molecules photobleached in single steps, while aggregated molecules photobleached in multiple steps (Fig. 2). We observed that 38% of spots are real single molecules, where 36% of spots are aggregated molecules. Only the sequences from the real single molecule spots will be used for analysis.

**Figure 2.**
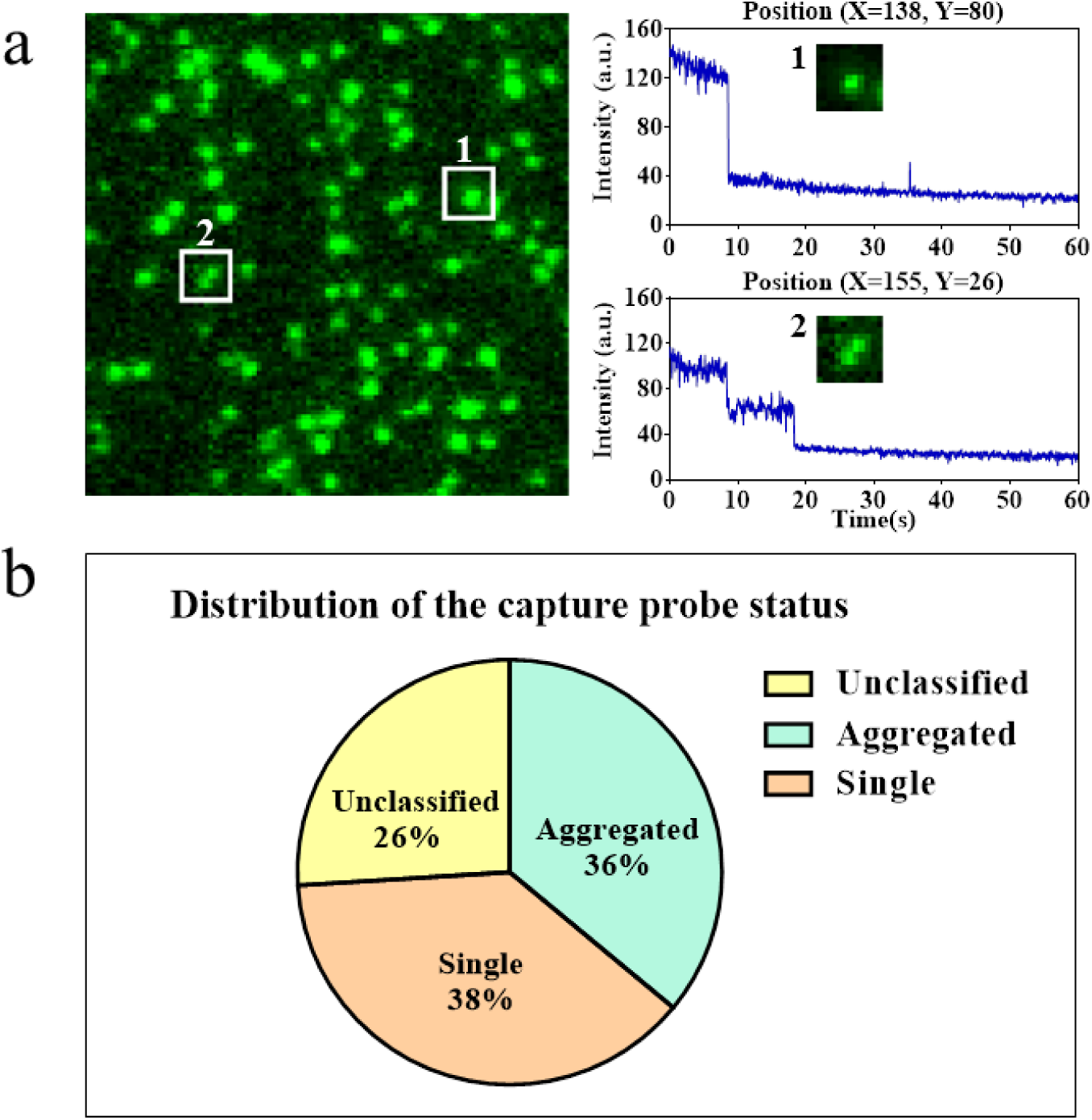
Quantifying the ratio of single molecules of capture probes. (a), the photobleaching of single molecules are in a single step. Here, single spot was traced and its intensity was recorded. A single-step photobleaching indicated that this spot was composed only one Cy3 molecule, i.e the spot #1. Spot #2 was composed of two molecules binding together and therefore displaying two steps of photobleaching. (b), The composition of single molecules, aggregated molecules and unclassified cases in one field of view.

### Targeted hybridization and sequencing

The EGFR, KRAS, BRAF genes were selected for sequencing in this studies. In particular, we aimed to sequence the 8 genetic variants that are related to drug response, including six point mutations and two short deletions (Table 1). Eight capture probe sequences were designed in the upstream of drug response related mutations. The capture probes are synthesized and anchored to the flow cell surface by a expoxy-NH2 bond. We synthesized two sets of target DNA templates for sequencing. The first set was wild type sequence and the second set contained mutations and short deletions (Table 1). Each target DNA template contained a Cy3 fluorescence dye at the 3’ end. Excitation of 3’ Cy3 fluorescent dyes was used to mark positions of annealed templates on the flow cell surfaces. Synthetic target DNA templates were hybridized to the flow cell with surface-attached capture probes (Fig 3a).

**Figure 3.**
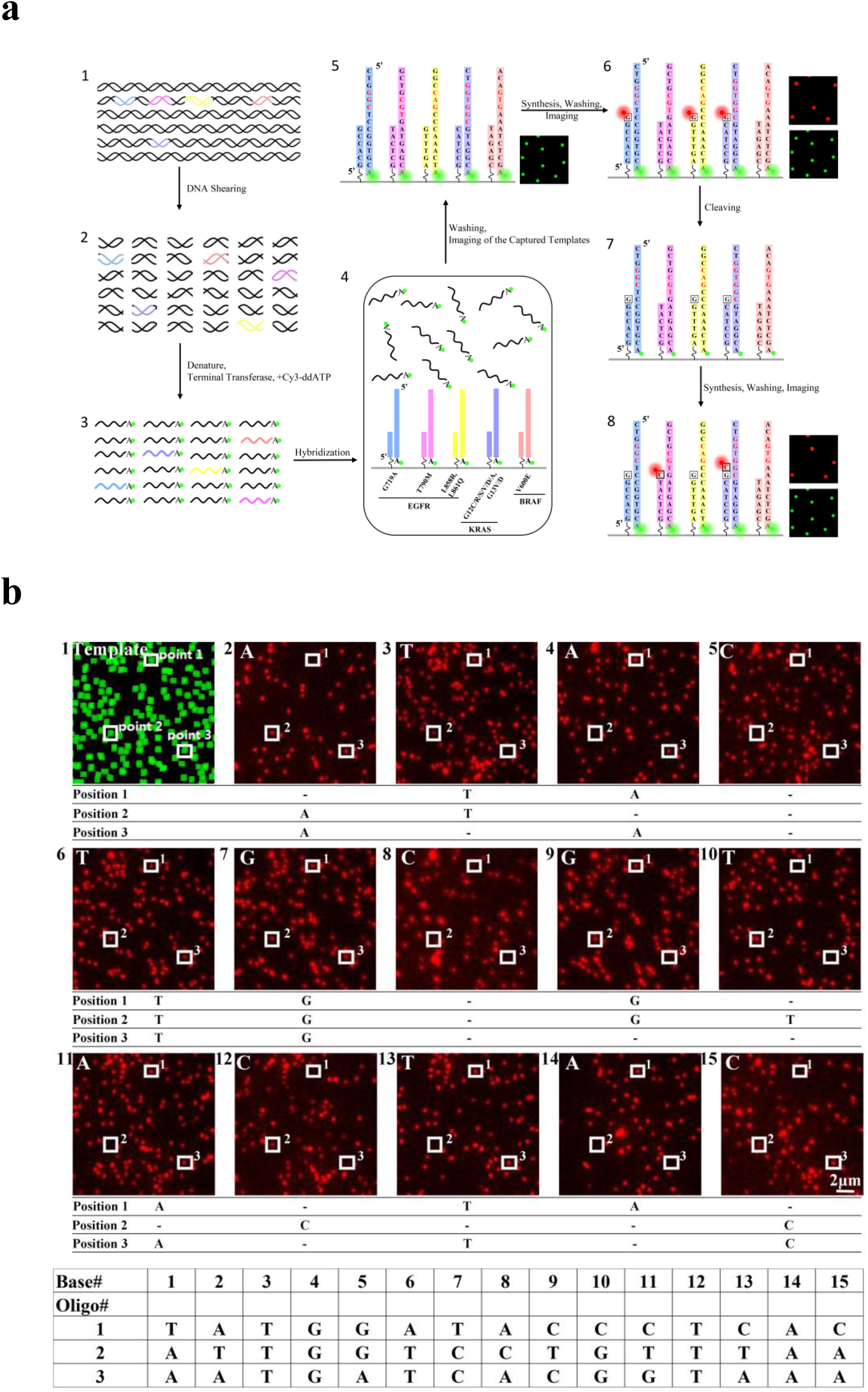
**(a)**, The sequencing procedure. DNA template with Cy3 attached at 3’ end was hybridized to the flow cell anchored with capture probes (step 1–4). The capture probes are designed complementary to the genes of interest. Unhybridized DNA templates were washed away. The green laser excited the Cy3 fluorescence dye to locate the position of target DNA templates. (step 5). One of four types of reversible terminators labeled with red fluorescence and polymerases mixture were added to the flow cell. The DNA molecule extended a base if the reversible terminator matched complementary to the next base in the DNA molecule. Unincorperated reversible terminator was washed out. The red laser excited the Atto647 fluorescence dyes of reversible terminators (step 6). The fluorescence dyes in the reversible terminators were cleaved and wash away (step 7). A new cycle of sequencing began (step 8). (b), Multiple sequencing cycles, imaging and base calling. We traced a part of one field of view in multiple sequencing cycles. In the beginning, the image of Cy3 green fluorescence dyes were used to locate the position of target templates. Three positions were circled out and were traced. In the first cycle, reversible terminators A (nucleotide analogs) were flowed in for reaction. Position 2 and 3 successfully incorporated a base. In the second cycle, the reversible terminators T were flowed in for reaction. Position 1 and 2 successfully incorporated a base. The sequencing continued and the sequence of DNA template extended. The sequence of each DNA template in position 1, 2 and 3 can be reconstructed.

**Table 1.**
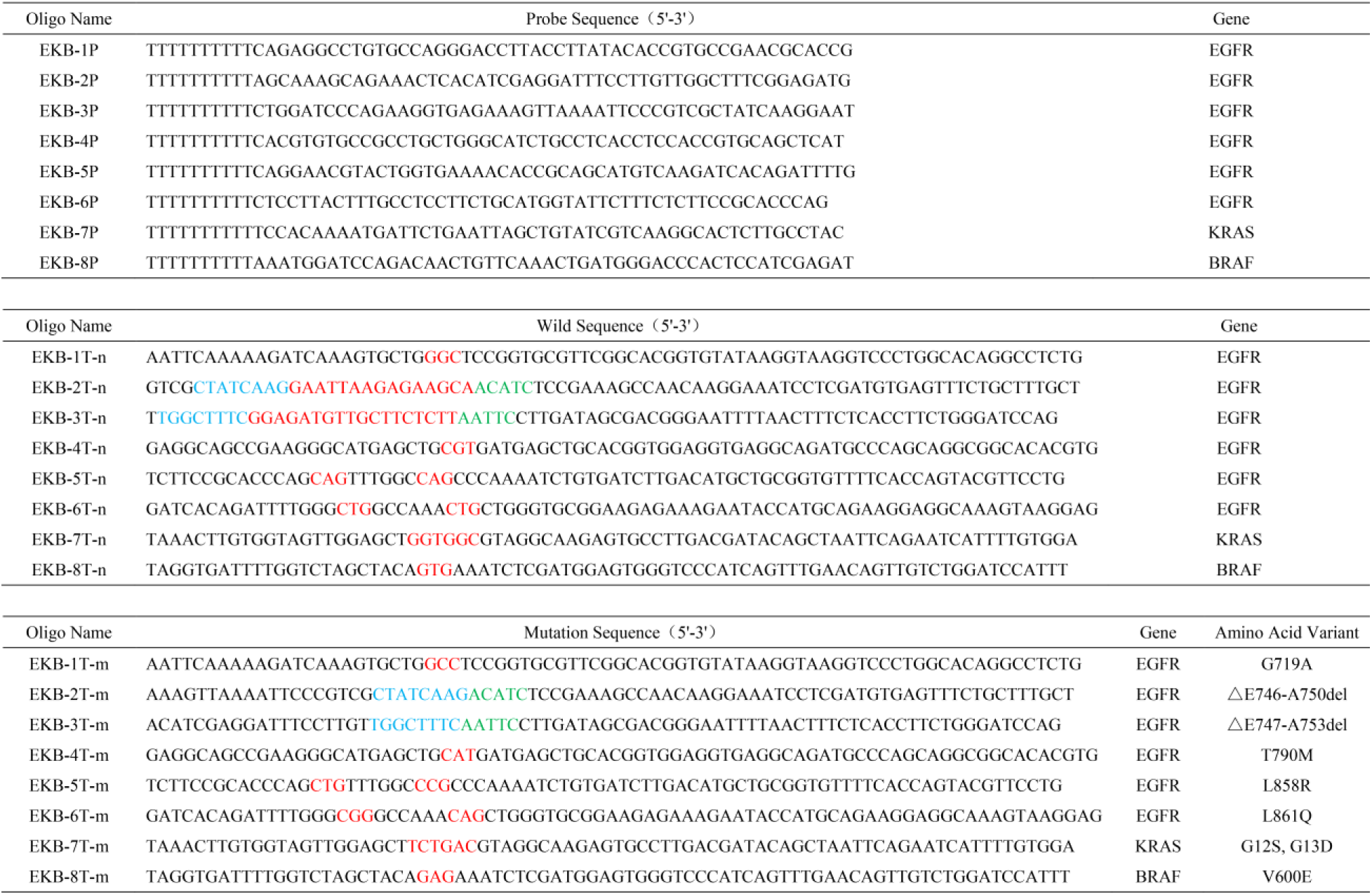
The capture probe and target DNA sequence information. The top block is the capture probe sequences we synthesized. The capture probes were designed to capture EGFR/KRAS/BRAF genes. The middle block is the target DNA sequence designed for testing. These sequences are design based on the wild type of EGFR/KRAS/BRAF genes. Nucleotide bases in red color are drug related mutation sites. The bottom block is the target DNA sequence designed based on the mutant type.

The sequencing reaction began with locating the target DNA templates, which are randomly hybridized to capture probes (Fig. 3a). The Cy3 fluorescent dyes attached to target DNA templates are excited by a 532nm green laser and the images were collected to locate the positions of target DNA templates. Then, disulfide linked Atto647N labeled reversible terminators and DNA polymerases were added to the flow cell. The reversible terminators were nucleotide analogs modified to contain a cleavable liner, which allowed only one reversible terminator to be incorporated into the DNA molecule at one time. The polymerase synthesis reaction was carried out at temperature 37°C, with one of four types of reversible terminators and necessary cofactors. Unincorporated reversible terminators were washed way. The Atto647N dyes are excited by a 640nm red laser in an optimized imaging buffer mixture with oxygen scavenging, free radical scavenging, and triplet quenching components. The images were processed using a custom written computer program to automatically locate the spot, determine image noise, and filter out false-positive spots. After imaging, the Atto647N fluorescence dyes were cleaved from the reversible terminators, and the system is ready for a second round of adding reversible terminators and polymerases. The sequencing cycle are repeated many times to achieve the desired length of read (Fig. 3b).

### Sequencing coverage depth

To demonstrate the performance of SMTS, we sequenced the wild-type EGFR/KRAS/BRAF DNA templates. The synthesized DNA templates were hybridized to the flow cell with surface-attached capture probes. We sequenced DNA for 19-30 cycles, which enable to cover all mutation/deletion loci. 300 fields of view were imaged for each cycle. In each field of view, there are approximate 2200-2500 reads on average. The sequencing reads were aligned to reference sequences with customized program of Smith-Waterman algorithm (Table 2). We observed that the coverage depth varies among different DNA templates (Figure 4a). The possible explanation is that the hybridization efficiency for DNA templates is sequence-dependent and the secondary structures that involve the target region can also affect hybridization efficiency. The average coverage depth was 1954-fold. Higher coverage depth can be achieved by capturing images for more fields of view.

**Figure 4.**
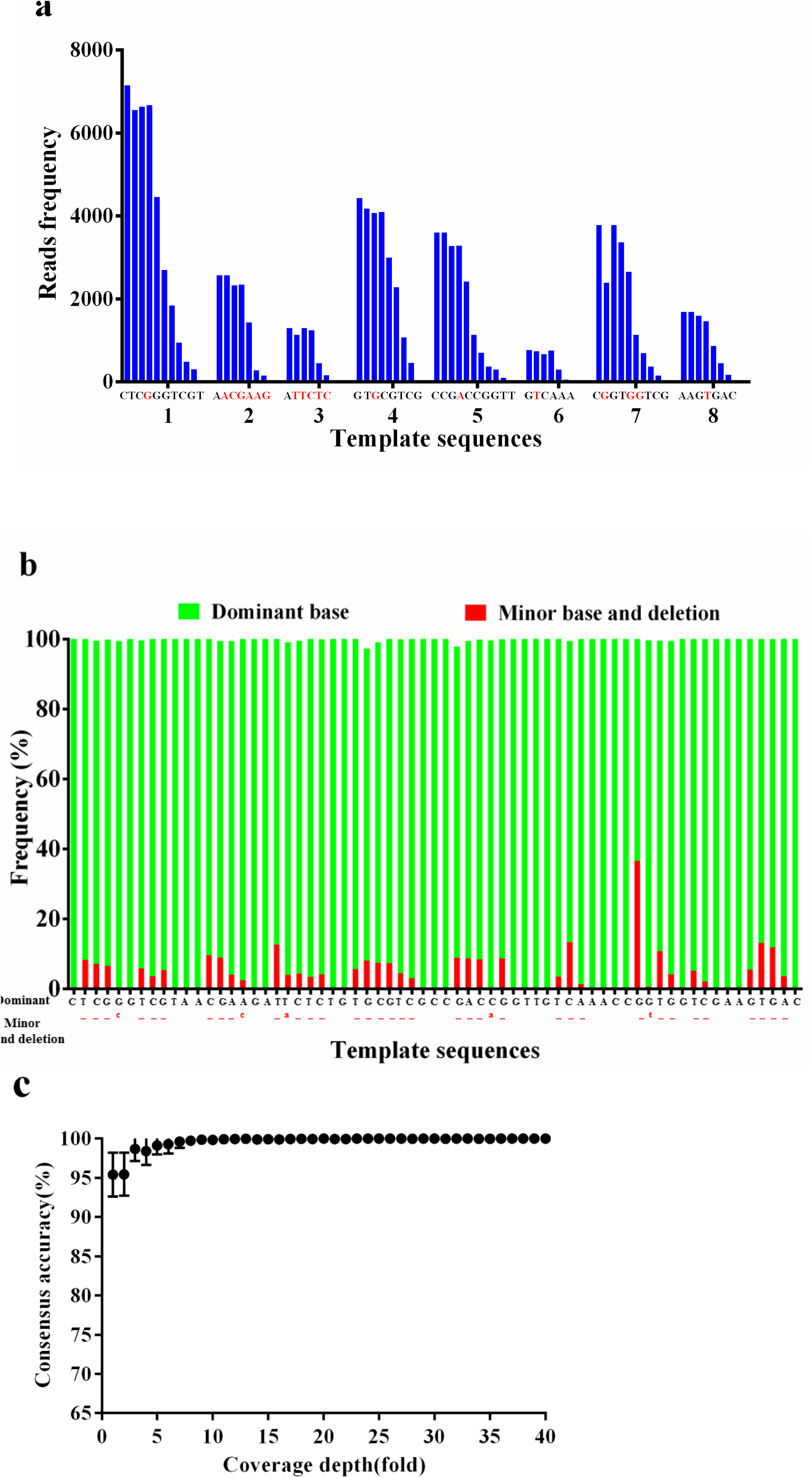
(a), the coverage per base. The frequently mutant position is in red color. Y-axis is the number 390 reads mapped to each position. (b), The dominant and minor base at the each position. (c), The consensus 391 accuracy increased with coverage depth. Sampling-subsampling was performed to simulate low coverage 392 situation.

**Table 2.**
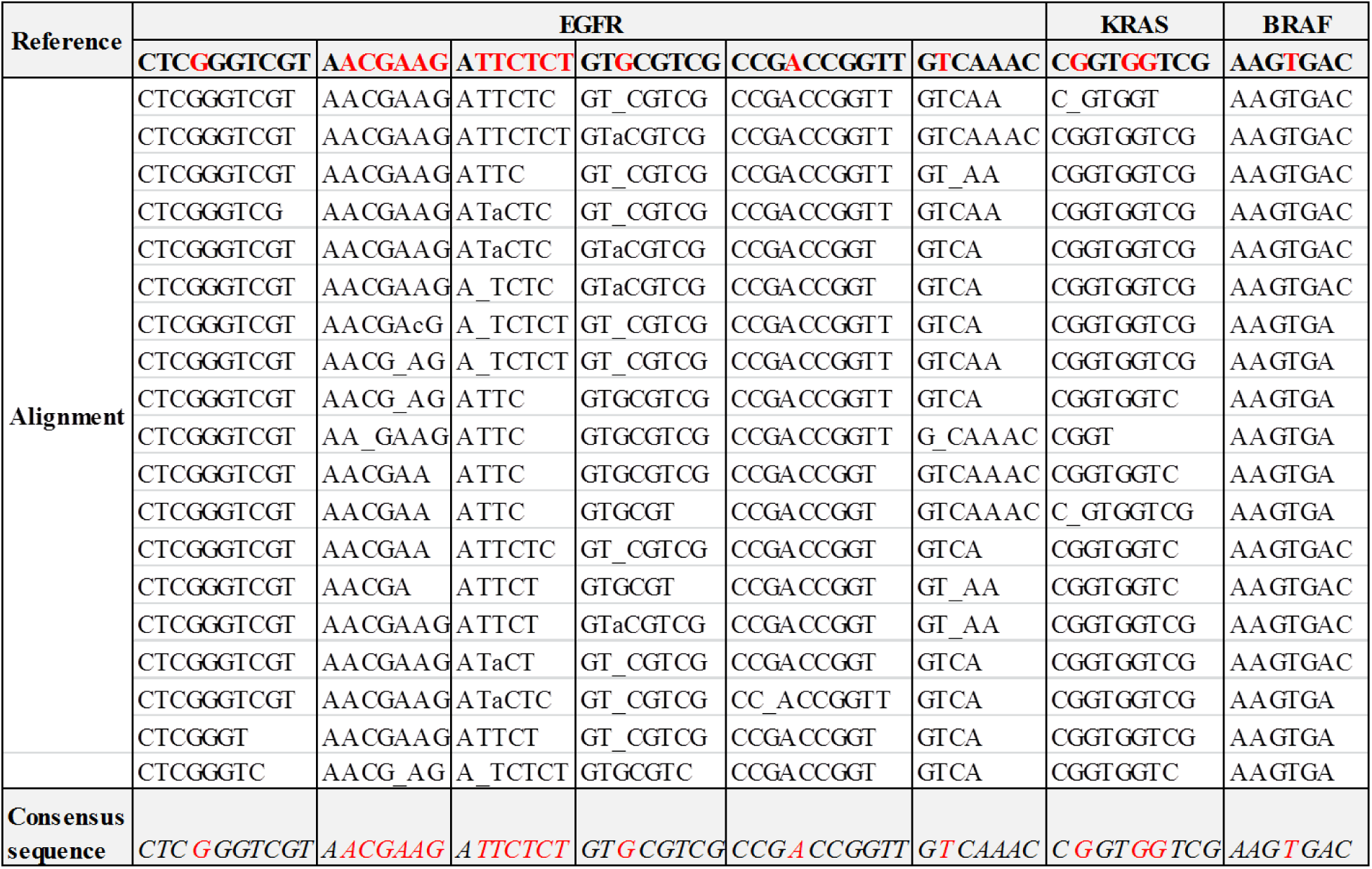
Sequencing alignment of raw reads. The top row is the reference sequence. Insertion errors were not shown in the alignment.

### Sequencing accuracy

The accuracy was calculated by comparing the reference sequences with the consensus sequences. Consensus sequences were calculated as the most frequent bases at each position in the sequence alignment (Table 2). By comparing the consensus sequence to the reference sequence base-by-base, the consensus sequence is 100% identical to the reference sequence in our four repeated experiments. We performed sampling-subsampling to the sequence data to get low-coverage data, and recalculated the consensus sequences at different coverage depth. If each base was covered only one time, which means the coverage depth is 1 fold, the accuracy was 95% on average. If each base was covered with 5 times or more on average, the consensus accuracy is approaching 100% accuracy (Fig. 4c). We performed multiple repeated experiments to estimate the errors in the raw sequencing data. The reads from each template were separately aligned to the DNA reference. Each position in the reference was mapped by multiple reads. The error rate of a position was the ratio of reads disagreeing with the reference divided by the total number of reads mapped to the reference. The overall error rate was an average of error rate of all positions. The error of raw sequencing reads was dominated by deletion (Fig. 4b). The substitution error is relatively small, in four repeated experiment, the average substitution rate is 0.52% per base (Fig. S6).

### Detecting low frequency of mutations

The wild type DNA was mixed with mutant type DNA at 10:1 and 97:3 ratios (Table 1). The DNA mixture was hybridized to the flow cell and sequenced. Each raw sequence read was aligned to reference sequences to determine whether it originated from wild type or mutant type DNA. As a control, we also sequenced pure wild type DNA with the same condition. We found that the percentage of mutant DNA detected in the DNA mixture was significantly higher than that in pure wild type DNA control (Fig. 5). In this experiment, SMTS can detect mutant sequences with frequency 3%.

**Figure 5.**
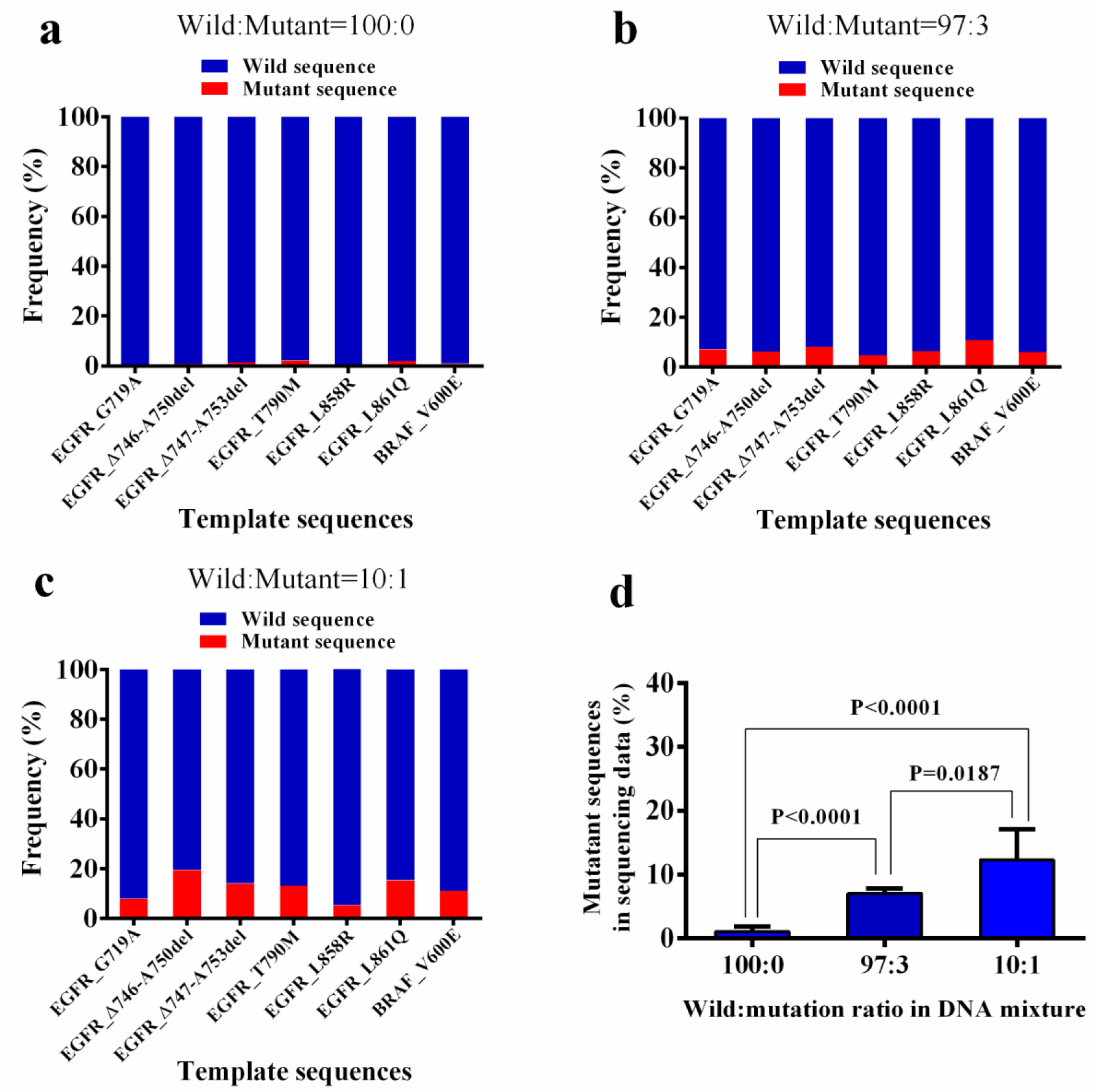
Detecting mutant sequences in a mixture. The wild type and mutant DNA were mixed at 100:0, 10:1 and 97:3 ratios. The mixed DNA was subjected to sequencing. Each sequence reads was aligned to the wild type and mutant type reference sequences and alignment scores were calculated. If the alignment score of wild type reference sequence was higher than that of mutant type reference sequence, the original sequence read was classified as wild type. Otherwise, it was classified as mutant type. The frequency of wild type and mutant type sequence reads are calculated for each reference. (a-c) The frequency of wild type and mutant type sequences calculated from the sequencing data. (d), The average of mutant sequences in sequencing data over all template sequences. P value is calculated by two-tailed Student T test.

## Discussion

We here demonstrated a method of capturing and sequencing DNA in a single step, which provides a much simpler approach to targeted sequencing. We have shown that the mutations and short deletions can be accurately detected at low frequency.

We have included several mutations of EGFR/KRAS/BRAF genes in this study. These mutations are actionable and can be therapeutically target. Somatic mutations in EGFR in exon 18, 19, 21 and the T790M point mutation in exon 20 are predictive of a clinical response to the EGFR tyrosine kinase inhibitor drugs gefitinib and erlotinib^9, 10^. Somatic mutations in KRAS (codons 12, 13) and BRAF (V600E) in colorectal cancer that predict poor prognosis and nonresponse to anti-EGFR antibodies. BRAF V600E is predictive of a positive response to the BRAF V600- specific inhibitor vemurafenib in melanoma^11^.

SMTS has several advantages over the more traditional Sanger sequencing and other NGS platforms commonly used for the detection of mutations. Firstly, there is little required in the way of sample preparation. Only sonication of the genomic DNA is needed. In the case of nucleic acids from sources such as FFPE or cfDNA, it is possible that even the sonication is not needed. Other high throughput sequencing technology such as Illumina requires days of labor work on sample preparation, which contains multiple steps such as sonication, end repairing, dA tailing, adaptor ligation, PCR amplification and target enrichment. Therefore, the SMTS technology has the potenial of reducing cost, turn-around time and the risk of errors in sample handling. Secondly, SMTS technology directly sequences original individual molecules, not PCR products. This should provide increased sensitivity for the detection of low prevalence mutations and avoid PCR biases^12^, which are essential features in the sequencing of a heterogeneous cancer sample^13^.

We observed that the coverage depth was not uniform among different positions. Some sequences appeared to be difficult to be sequenced. The uniformity of coverage could be improved by carefully designing the capture probes, in particularly, to avoid the secondary structure. We also observed that only one third of fluorescence spots were from single molecules. Under the random attachment scenario described in this study, a large portion of spots came from two or more molecules binding closer than the diffraction limited resolution of the system. The ratio of single molecules could be increased by optimizing the hybridization condition and/or controlling the density of capture probes. The overall error rate of raw sequences was still significant ^14^. To reduce the error rate, we need to further optimize the chemical reaction conditions for incorporating reversible terminators and cleaving the fluorescence dye after imaging. Meanwhile, by modeling the error profiling, a better base calling algorithm could be developed. The four reversible terminators (A, T, C and G) used in current study were labeled with the same fluorescence dye. In future, we can modify the reversible terminators and label each of four nucleotides with unique fluorescence dyes^15^. By doing so, the speed and accuracy will be improved.

For the foreseeable future, the high cost and complexity of data analysis will limit the application of whole-genome sequencing for the detection of mutations in a clinical setting. Targeted resequencing of areas of interest will therefore remain key to determining mutational status. SMTS is a stride forward in putting this into practice. Although currently only a few loci of a few genes are screened, there is clearly scope for the creation of multi-gene capture arrays, allowing large numbers of loci to be analyzed rapidly and cost-effectively with low DNA input requirements. The single-step capturing and sequencing whole exome is also possible in future. In its simplicity, this approach provides an opportunity to truly begin integrating the vast quantity of genomic data generated in this next-generation era with clinical practice.

## Methods

### Optical setup

A custom-engineered sequencer prototype contained a Total Internal Reflection Fluorescence (TIRF) microscope with 60X oil objective (Nikon Ti-E, Japan), EMCCD camera with a resolution of 512X512 (Andor, Belfasst, UK) and 2 color laser powers, 532nm (100mW) and 640nm (100mW). A motorized stage (ASI, Eugene, OR) was installed on the TIRF microscope to hold and control the motion of the flow cell (Bioptechs, Bulter, PA) during sequencing. The heater (Bioptechs, Bulter, PA) for both flow cell and objective was installed and can maintain the temperature in chamber of the flow cell at 37°C.

### Flow cell and liquid handing

The FCS2 flow cell contained the chemical functionalized coverslip with epoxy layer (Schott, Jena, Germany), 0.175mm thick and 40mm in diameter. A gasket was assembled between the coverslip and an aqueduct slide which forms the chamber (3mm X 23 mm X 0.25 mm) for chemical reaction. The sandwiched structure part was fixed by a top with stainless steel tube inside (inlet port and outlet port) and metal base. A Titan EZ valve with 12 channels (IDEX Health & Science, Oak Harbor, WA, USA) was connected between the inlet of the flow cell and sequencing reagents. The outlet of the flow cell was connected with a syringe pump (Tecan, Männedorf, Swiss) to drive the fluidic in the system by suction.

### Surface chemistry

Synthesized capture probes (oligonucleotides) were covalently coupled to the epoxy coated coverslip surface. The capture probes were firstly incubated at 95°C, then the coverslip was immerged into a capture probe solution at 1 nM in 150mM K_2_HPO4, pH 8.5 at 37°C for 2 hours. Then the coverslip was rinsed by 3X SSC with 0.1% Triton X-100 and 3X SSC, 150mM K_2_HPO4, pH 8.5 in sequence.

### Imaging processing

Images are processed using a custom written spot localization algorithm (Fig. S4). Firstly, stage drifts between different imaging cycles were corrected by calculating the peak position of two images by Phase-Only Correlation (POC) function. After correcting all cycles with the corresponding first cycle, the corrected images were convolved with a Gaussian kernel. The correlation images were then subjected to the threshold determined by the noise measurement on those images. All contiguous groups of pixels above the threshold were grouped as spots. After that, each spot was fitted with a Gaussian function. This step allowed an accurate determination of the centroid position for single molecules and both members of closely standing molecule pairs. At the same time, clusters of three or more molecules were filtered out. A spot that appeared twice at a same time point but under different wavelength lasers was considered as a base incorporation event. Thus, the spot was renamed as an incorporation spot and marked on the incorporation image. A set of incorporation spot centroids falling within a 1.6 pixel radius is called a “track”. Comparing with the order of adding reversible terminators, these “tracks” were converted to the final sequences on the position of each incorporation spot.

### Target template of EGFR/KRAS/BRAF

Eight mutation sites in three genes (EGFR, KRAS and BRAF) were covered by the target templates, including six point mutations (G719A in EGFR exon 18, T790M in EGFR exon 20, L858R and L861Q in EGFR exon 21, G12S and G13D in KRAS exon 2 and V600E in BRAF exon 15) and two short deletions (ΔE746-A750 deletions and ΔE747-A753 deletions in EGFR exon 19). We designed two target sequences for each genetic variant, which are wild type and mutant type. The length of each target template was 70 bp, with a Cy3 fluorescence dye attached to the 3’ end. Synthetic target templates were hybridized with capture probes attached on the surface of flow cell according to complementary matching principle.

### Capture probe design

A 60nt capture probe sequence with 10 dT bases and an amine labeled 5ʹ end was designed according to the upstream gene sequence of mutation sites. The 50nt target-specific sequence at the 3ʹ end of capture probe sequence were designed according to the program BatchPrimer3, with specified conditions: 20%–80% GC and Tm’s >65 °C. Capture probes and target templates were synthesized by Sangon Biotech(Shanghai).

### Reversible terminators

The modified reversible terminators are composed of nucleotide triphosphates, modified with a detectable label (Atto647N) by disulfide linker and an inhibitor group(SeqLL, Woburn, MA, USA). The inhibitor region has multiple negative charged groups (carboxyl group) allowing incorporation of one nucleotide into the DNA duplex while prohibiting the second or third or more nucleotide incorporation. The detectable label and inhibitor group were cleavable.

### Sequencing cycle

The coverslip was incubated in synthesized templates labeled with Cy3 solution at 5nM in 3X SSC, pH7, at 55 °C for 2 hours to form a DNA duplex. Then the surface was rinsed with 150mM HEPES, 1X SSC and 0.1% SDS, followed by 150mM HEPES and 150mM NaCl. Finally the coverslip was assembled into the follow cell.

The sequencing process was controlled automatically by the fluidic system. Two different types of reagents containing nine pre-prepared reagents were used and stored at different temperatures. One type is the chemical or biochemical reaction reagents, including four nucleotide (dNTP-Atto647N) and DNA polymerase mixtures, cleavage reagent (TCEP, 50mM), cap reagent (50mM idoacetamide), and imaging buffer (50mM Trolox, 20mM glucose and 5mM glucose oxidase in HEPES buffer) stored at 4°C. The other is rinse buffer including rinse buffer 1 (150mM HEPES, 1X SSC and 0.1% SDS, pH 7.0) and rinse buffer 2 (150mM HEPES and 150mM NaCl, pH 7.0) stored at room temperature.

First, 0.25 μM reversible terminators (one of G, C, T and A) and 20nM polymerase mixture was introduced into the flow cell, incubated for 4 minutes at 37 °C and washed out by rinse buffer1 and 2. Then imaging buffer (50mM Trolox, 20mM glucose and 5mM glucose oxidase in HEPES buffer) was pumped in the flow cell. Then, the images of 300FOVs were taken. Typically, 4 exposures of 0.1 second were taken in each field of view (FOV, 54.6μm ×54.6μm). After imaging, the flow cell was washed by rinse buffer. The cleave reagent was introduced into the flow cell and reacted for 5 minutes flowed by the cap reagent under reaction for another 5 minutes. Finally the flow cell was washed by rinse buffer and finished the first cycle of sequencing. The sequencing cycle was repeated with the same procedure, except changing the reversible terminators. In this paper, the terminators were added into the system as the repeated order of G, C, T, A.

### Bioinformatics

Quality control on the sequence reads was first performed. Firstly, reads with length less than 5 bases were filtered out. Then, sequencing reads that appeared less than 4 times were filtered out. Secondly, sequencing reads that could not be aligned to reference sequences are not included for further analysis.

The alignment described above was performed with Smith-Waterman algorithm, which performs local sequence alignment. By using a custom definition scoring system (which included the substitution matrix and the gap-scoring scheme), the chosen algorithm could guarantee identification find of the optimal local alignment. In this setup, the penalty for a deletion in a read was −1, for an insertion −1, for a match 2, and for a substitution −2.

## Acknowledgement

This work was financially supported by the Shenzhen Science and Technology Innovation Commission (#Pu20150387 and #CYZZ2014110845956), and in part by Human resources and social security bureau of Shenzhen City (#2015069). We thank Dr. J. William Efcavitch and Dr. Yilin Wang for helping on designing and performing experiments. We also thank Azco and SeqLL for providing help on reagents.

